# Large-scale *in silico* mutagenesis experiments reveal optimization of genetic code and codon usage for protein mutational robustness

**DOI:** 10.1101/2020.02.05.935809

**Authors:** Martin Schwersensky, Marianne Rooman, Fabrizio Pucci

## Abstract

The question of how natural evolution acts on DNA and protein sequences to ensure mutational robustness and evolvability has been asked for decades without definitive answer. We tackled this issue through a structurome-scale computational investigation, in which we estimated the change in folding free energy upon all possible single-site mutations introduced in more than 20,000 protein structures. The validity of our results are supported by a very good agreement with experimental mutagenesis data. At the amino acid level, we found the protein surface to be more robust to mutations than the core, in a protein length-dependent manner. About 4% of all mutations were shown to be stabilizing, and a majority of mutations on the surface and in the core to be neutral and destabilizing, respectively. At the nucleobase level, single base substitutions were shown to yield on average less destabilizing amino acid mutations than multiple base substitutions. More precisely, the smallest average destabilization occurs for substitutions of base III in the codon, followed by base I, bases I+III, and base II. This ranking highly anticorrelates with the frequency of codon-anticodon mispairing, and suggests that the standard genetic code is optimized more to limit translation errors than the impact of random mutations. Moreover, the codon usage also appears to be optimized for minimizing the errors at the protein level, especially for surface residues that evolve faster and have therefore been under stronger selection, and for biased codons, suggesting that the codon usage bias also partly aims to optimize protein mutational robustness.

## 1 Introduction

Amino acid substitutions can have different impacts on protein fitness. Some have highly destabilizing effects, thus causing the loss of structure and/or function. Others lead to the emergence of new functions, although with a very low frequency of about 10^−9^ per site, thus driving functional evolution [1]. But the large majority of amino acid substitutions are neutral with respect to protein fitness [46].

Two concepts play a central role in these matters: mutational robustness, which refers to the capacity to tolerate mutations without changing the molecular and/or organism’s phenotype, and evolvability, which is defined as the capacity of proteins of acquiring new functions, hence allowing them to adapt to modifications in the environment.

Despite recent advances, the role of the evolutionary mechanisms in the complex interplay between the optimization of these two fundamental but sometimes conflicting characteristics is still a major issue in molecular evolution and protein biophysics [13, 53, 14, 64, 66, 73, 69, 70]. A wide variety of disciplines, from synthetic biology to protein design, would definitely benefit from a better understanding of these mechanisms and from the ability of accurately predicting the future evolutionary processes from the analysis of the past [51].

Mutational robustness and evolvability can be viewed as two sides of the same coin, which drive evolution in an entangled way. On the one hand, physical principles are expected to favor proteins with a high degree of stability, while on the other hand the selection for function imposes opposite constraints in targeted regions, such as the presence of amino acids carrying specific chemical moieties or a required degree of structural flexibility. Once the functional criteria are satisfied, mutational robustness ensures better tolerance of random mutations in non-functional regions and thus confers an evolutionary advantage [12, 10]. Note, however, that too high tolerance to mutations can also prevent necessary adaptation to environmental changes [26].

Results obtained from experimental analyses and theoretical models of population genetics suggest that mutational robustness is favored or disfavored, and impedes or facilitates adaptative evolution, according to the polymorphicity and size of the population, the mutation rate, and the fitness landscape [72, 14, 15, 26].

To further shed light on these challenging issues, we performed an extensive *in silico* mutagenesis study, in which we computed the change in protein thermodynamic stability caused by all single point mutations inserted in the structurome, defined as the ensemble of all protein structures available in the Protein Data Bank [9]. To support our analyses, we also included available experimental data on stability changes and fitness.

The first question that we investigated in detail on the basis of these large-scale computations is how the mutational robustness is influenced by some protein characteristics, such as protein length and residue solvent accessibility.

A second question concerns the relation between the mutational robustness and the standard genetic code (SGC). It has been shown that this code has evolved to minimize the costs of amino acid replacements. Indeed, from the observation of the SGC table (Fig. S1), we immediately see that amino acids that share similar biophysical characteristics tend to be encoded in codons that differ by only a single base. However, a long and controversial debate regards the level of optimality that SGC has reached. [40, 31, 39, 36, 23, 38, 76].

We also retrieved the nucleobase sequence of the whole structurome and this allowed us to investigate the relation between the mutational robustness, the codon choice and the codon usage bias. Indeed, the degeneracy of the genetic code introduces some variability into protein encoding in nucleobase sequences, which opens alternative pathways in the evolutionary landscape that are likely to allow, *e.g.*, the minimization of translational errors and an effective increase of protein mutational robustness [16, 4, 5]. Codons are selected for other reasons too, such as the matching of tRNA abundance and the mRNA stability for improved translation efficiency [5, 43, 44, 2].

## 2 Results and Discussion

The central question addressed here concerns the protein robustness against mutations, its dependence on various parameters at the codon, residue and protein levels, and its link with evolutionary rates.

With this objective in mind, we estimated with the PoPMuSiC^*sym*^ algorithm [59, 60] the change in folding free energy (ΔΔ*G*) for all single-site mutations in the non-redundant set 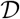 of protein X-ray structures representing the protein structurome, as described in Methods. In parallel, we considered the smaller ensembles of experimentally measured ΔΔ*G* values and fitness scores. These three sets of mutations, that we call 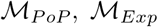 and 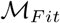 contain about 4 × 10^5^, 2.6 × 10^3^ and 10^4^ mutations, respectively.

Most of the natural amino acid mutations are the result of a single base substitution (SBS) in the codon, as the evolutionary probability to have simultaneously two or three base substitutions is small. However, we would like to point out that only a subset of all possible amino acid mutations can be obtained through SBSs. We call such amino acid mutations *μ*SBS, and limit ourselves to this subset unless otherwise stated. The amino acid mutations that result from multiple base substitutions (MBS) are called *μ*MBSs.

### 2.1 Relative solvent accessibility

We started by analyzing the effect of the relative solvent accessibility (RSA) of the mutated residues on the mutational robustness. This effect is clearly visible in Figs 1.a-b: the ΔΔ*G* distribution of *μ*SBSs is much more spread out and shifted towards destabilizing mutations for core residues than for surface residues. Random mutations are thus on average much more destabilizing when introduced in the core, where close packing and specific interactions tend to impede changes in residue size and physicochemical properties. In contrast, surface residues are more robust to mutations than core residues, in the sense that they have a smaller impact on the thermodynamic stability.

**Figure 1:**
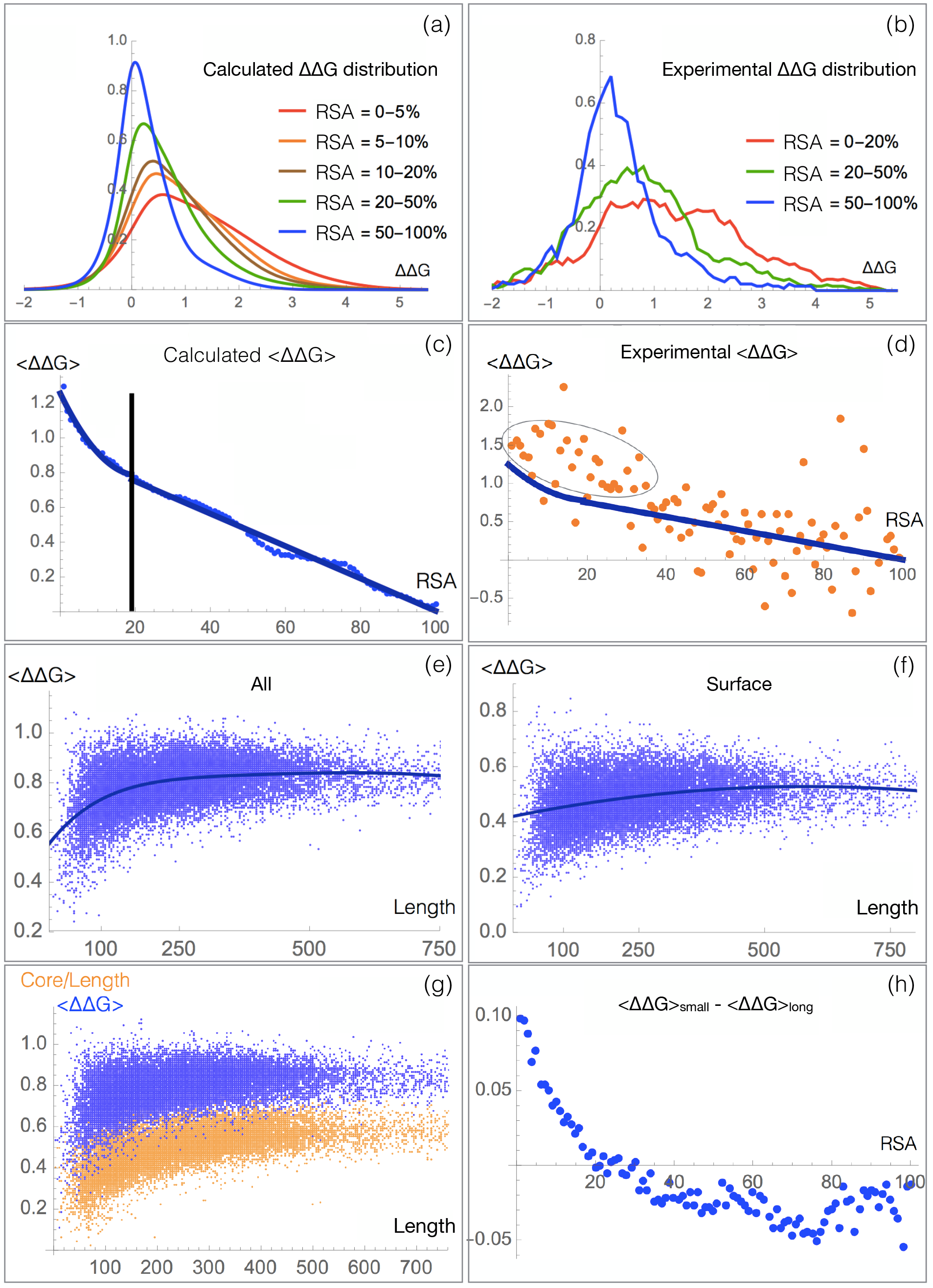
Influence of the protein length and of the mutated residues’ RSA (in %) on the mutational robustness, evaluated from the ΔΔ*G* values (in kcal/mol) of *μ*SBSs from the sets 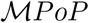 (a,c,e-h) and 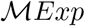 (b,d). (a-b) ΔΔ*G* distribution for different RSA ranges; (c-d) Mean ΔΔ*G* per RSA bin as a function of the RSA; the chosen bin width is equal to 1%; (e) Mean ΔΔ*G* per protein as a function of the protein length for all residues and (f) for surface residues (RSA≤20%). (g) Mean ΔΔ*G* per protein as a function of protein length (blue points) and protein core to length ratio, defined as the number of residues in the core over the number of residues in the protein (orange points); (h) Difference between the mean ΔΔ*G* per protein and per RSA bin of long proteins (*L* >200 residues) and short proteins (*L* ≤200 residues) as a function of RSA.

It has to be stressed that these results are almost identical whether using the set of computed or experimental ΔΔ*G*s from 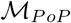 and 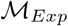 (*cf.* Figs 1.a and b). This supports the validity and accuracy of PoPMuSiC^*sym*^’s ΔΔ*G* predictions.

We also found that the relationship between RSA and ΔΔ*G* values is linear above an RSA threshold of about 20% and non-linear below this threshold, where the curve is well fitted by a second-degree polynomial function (Fig. 1.c-d). Again, the same trend is observed for the computed and experimental mutations of 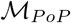 and 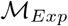, with an even stronger deviation from linearity at small RSA values for the latter; note that the number of mutations in 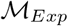 is low, which explains the noisy behavior.

### 2.2 Protein length

The effects of residue RSA and protein size on the mutational robustness are entangled. Indeed, mutations of residues located in the core, which have a low RSA, have on average a larger impact on stability than surface mutations, which have a large RSA. As a consequence, proteins of different sizes, which have different core to surface ratios, appear to have different tolerances to mutation [33].

The dependence of mutational robustness on protein length *L* is shown in Fig. 1.e. On average, shorter proteins that have a smaller core to surface ratio are more robust than longer proteins for which this ratio is larger. Above about 400 residues, the robustness remains roughly constant. Such large proteins are usually multi-domain proteins, which implies that the core to surface ratio does not increase any more.

To gain insights into this effect, we computed the *L*-dependence separately for core and surface residues. We found that shorter proteins tend to have a less robust core and a more robust surface than larger proteins, as shown in Figs 1.e-f and S2.

The former observation can be attributed to the larger compactness and hydrophobicity of the core of short proteins, which is therefore less able to accommodate mutations. We indeed checked that the core becomes less and less hydrophobic as the protein size increases (Fig. S3). In fact, the increase in core to surface ratio is compensated up to a certain level by variations in the amino acid composition. However, this compensation is far from perfect, and the core of small proteins is definitely more hydrophobic than that of large proteins [22]. For example, the hydrophobic residues (Val, Ile, Leu, Phe) represent about 45% of buried residues in proteins of *L* ≤ 200, 41% for medium-size proteins (200 < L ≤ 200) residues and only about 37% in larger proteins (400 < *L*).

The second observation, *i.e.* the higher mutational robustness of the surface of small proteins compared to the surface of longer proteins, can be explained by the larger fraction of functional residues. These residues are known to be well optimized for function but poorly for stability [20, 37]. Therefore, their substitutions are likely to be stabilizing, which confers a higher mutational robustness to the surface of small proteins.

Finally note that the mean ΔΔ*G* per protein (〈ΔΔ*G*〉) is, on average, proportional to the fraction of residues in the core, as seen in Fig. 1.g. This follows from the fact that core mutations have much larger ΔΔ*G* values on average than surface mutations, and their effect thus dominates when computing the mean.

### 2.3 Evolutionary rate

We compared the mutational robustness analyzed in the previous sections with the evolutionary rate that has been estimated in a series of papers on the basis of sequence evolution models [35, 62, 78, 34, 63]. These two quantities are expected to be related given that stability is known to be one of the major factors contributing to the evolutionary pressure [77, 30, 29].

The dependence of the evolutionary rate on RSA was investigated in [35, 62]. A larger rate was found for surface than for core residues. This is in agreement with our findings of a larger mutational robustness. In brief, surface residues, whose mutations have on average smaller effects on protein stability, evolve faster than buried residues.

However, while the relationship between RSA and evolutionary rate appears to be linear [35, 62], the relationship between RSA and mutational robustness is shown to be linear only for RSA values larger than 20% (Figs 1.c-d). This could suggests either that our models or those of [35, 62] are not totally accurate, or that the correlation between mutational robustness and evolutionary rate is linear solely for surface residues and becomes non-linear for core residues, where the robustness decreases more than the evolutionary rate.

Finally, the RSA-evolutionary rate regression line has a larger slope for large proteins than for small proteins [35, 62]: the rate difference is almost zero for core residues and positive for surface residues. This means that surface residues from large proteins evolve faster than those from small proteins, whereas almost no difference is observed for core residues.

This can seem in contradiction with our results of small proteins having a more robust surface and a less robust core than large proteins (Fig. 1.h).

This is actually not the case. Rather, we run here up against the limits of the correlation between evolutionary rate and mutational robustness: small proteins have a more robust and slower evolving surface than large proteins, and a less robust and equally evolving core. The interpretation of this difference lies in the fact that a significant proportion of surface residues are functional, especially in small proteins. These functional residues increase the robustness by lowering the 〈ΔΔ*G*〉 as they are not optimized for stability, and decrease the evolutionary rate as many mutations render the protein non-functional.

### 2.4 Experimental fitness

To compare the computed mutational robustness with experimental fitness measures, we subdivided the mutations into stabilizing, neutral and destabilizing, using the free energy thresholds: ΔΔ*G* < −0.5 kcal/mol, −0.5 kcal/mol≤ ΔΔ*G* ≤ 0.5 kcal/mol and 0.5 kcal/mol< ΔΔ*G*, respectively.

With these definitions, the fractions of destabilizing, neutral and stabilizing *μ*SBSs from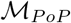 are (68%, 28%, 4%) in the core, (41%, 55%, 4%) on the surface and to (55%, 41%, 4%) overall (Fig. 2, and Table S1 for more detailed RSA dependence).

**Figure 2:**
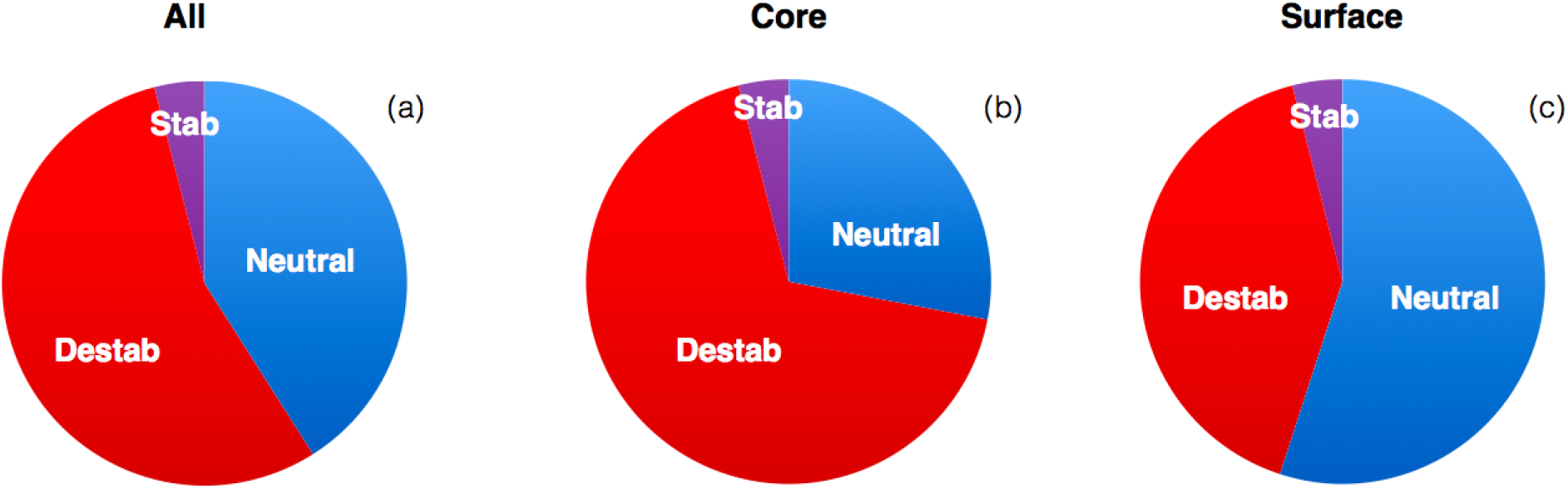
Fraction of stabilizing *μ*SBSs (ΔΔ*G* < −0.5 kcal/mol), neutral *μ*SBSs (−0.5≤ ΔΔ*G* ≤ 0.5 kcal/mol) and destabilizing *μ*SBSs (ΔΔ*G* > 0.5 kcal/mol) in 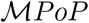 for: (a) all residues; (b) core residues (RSA≤ 20%); (c) surface residues (RSA> 20%).

Note that, in the set of experimental *μ*SBSs of 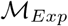, the fraction of stabilizing mutations is slightly higher (about 10 to 12 %, according to whether they are introduced in the core or at the surface). This is not surprising as these mutations are non-random; they are engineered and biased toward stabilizing mutations.

The fraction of stabilizing mutations obtained via a single base substitution is thus constant and equal to 4% of the total number of mutations both in the core and on the surface. In contrast, destabilizing *μ*SBSs dominate in the core and neutral *μ*SBSs dominate on the surface. Of course, the precise fractions of stabilizing, neutral and destabilizing mutations depend on the somewhat arbitrary threshold energy values of −0.5 and +0.5 kcal/mol.

We compared these results with experimentally characterized fitness values of random mutations, taken from three different studies and grouped in 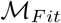 (Table 1). Note that the concept of fitness is not precisely defined and depends on the experimental setup used to characterize it. Stability is for sure a major factor [77], but fitness contains also other factors, related to, *e.g.*, protein expression, solubility and function.

**Table 1:** Comparison between mutational robustness and fitness: computed fraction of destabilizing, neutral and stabilizing *μ*SBSs from 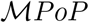 and experimentally characterized fraction of deleterious, neutral and advantageous mutations. The fitness thresholds for defining the mutation phenotypes are chosen by the authors for mutations in [54]; for the other sets of experimental mutations: deleterious if the fitness is lower than the mean of loss-of-function and wild-type scores, neutral if the fitness is between that threshold and 1.25 times the wild-type score, and advantageous otherwise.

| Mutation set | Destabilizing | Neutral | Stabilizing | Reference |
| --- | --- | --- | --- | --- |
| $\mathcal{M}_{PoP}$ | 55% | 41% | 4% | this paper |
| Mutations in | Deleterious | Neutral | Advantageous | Reference |
| AraC/D/E | 53% | 43% | 4% | [54] |
| UBE2I/SUMO1/CALM1/TPK1 | 51% | 44% | 5% | [75] |
| TEM-1 | 37% | 59% | 4% | [45] |

The first study involves about 150 mutations inserted in three proteins (the transcription factor AraC, the enzyme AraD and the transporter AraE) [54]. Among these mutations, the number of deleterious, neutral and advantageous mutations were found to be equal to 53%, 43% and 4% on average, with some differences between the three tested proteins. These values are close to the fractions of destabilizing, neutral and stabilizing mutations that we predicted for the full structurome.

A second experimental investigation used deep mutagenesis scanning to investigate about 13,000 mutations in four proteins (SUMO E2 conjugase, a small ubiquitin-like modifier, thiamin pyrophosphokinase and calmodulin). The percentage of deleterious (51%), neutral (44%) and advantageous (5%) mutations [75] also fits very well with our predictions.

The third series of experimental results concerns the mutational landscape of TEM-1 *β*-lactamase [45]. In this case, a bigger fraction of neutral than of destabilizing mutations was found. This could suggest that the activity of this enzyme is particularly well optimized as already observed in [45].

### 2.5 Similarity matrices

Similarity matrices, such as the series of BLOSUM matrices [42], are commonly used in sequence alignment methods to account for the similarity between the 20 amino acids and the ease with which they are mutated into each other. They are derived from multiple sequence alignments of homologous proteins and thus reflect both the physicochemical similarity of the substituted amino acids, the evolutionary mechanisms acting on protein sequences, and the structure of the genetic code.

We expected a certain correlation between BLOSUM scores and mutational robustness, as they share stability as one of their main ingredients. Moreover, BLOSUM and fitness scores have been shown to correlate in the mutational landscape of TEM-1 *β*-lactamase [45].

We focused here on mutational robustness rather than fitness, expanded the analysis to the ensemble of all *μ*SBSs in the structurome set 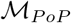, and computed the ΔΔ*G* distribution as a function of the BLOSUM scores. We considered for that purpose the commonly used BLOSUM62 matrix.

We clearly observe a strong correlation between the mean ΔΔ*G* and the BLOSUM62 score, with a linear correlation coefficient as high as *r*=−0.97. As shown in Table 2, the substitutions that are the most likely to occur during natural evolution are mostly neutral for stability and only a small fraction is destabilizing. The picture is completely reversed for the substitutions that are less likely to occur. Indeed these substitutions impact on average quite strongly on protein stability, while only a very small fraction is neutral. Interestingly, the fraction of stabilizing mutations is almost constant, between 3 and 5%, except for mutations between very similar amino acids where it drops to 1%.

**Table 2:** Mean ΔΔ*G* (in kcal/mol) of all *μ*SBSs in 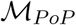 as a function of the BLOSUM62 class. Positive BLOSUM scores indicate more likely amino acid substitutions and negative scores, less likely substitutions. The fraction of stabilizing (Stab), neutral (Neut) and destabilizing (Dest) substitutions in each class is also reported.

| BLOSUM | $\langle \Delta\Delta G \rangle$ | Stab | Neut | Dest |
| --- | --- | --- | --- | --- |
| -4 | 1.58 | 4% | 18% | 78% |
| -3 | 1.15 | 5% | 29% | 66% |
| -2 | 1.11 | 4% | 27% | 69% |
| -1 | 0.83 | 4% | 34% | 62% |
| 0 | 0.56 | 5% | 46% | 49% |
| 1 | 0.33 | 3% | 65% | 32% |
| 2 | 0.28 | 3% | 71% | 26% |
| 3 | 0.25 | 1% | 78% | 21% |

The relation between mutational robustness and BLOSUM scores is clearly seen in Fig. 3: the ΔΔ*G* distribution extends more and more toward positive values – *i.e.* toward destabilizing mutations – when the BLOSUM62 score decreases.

**Figure 3:**
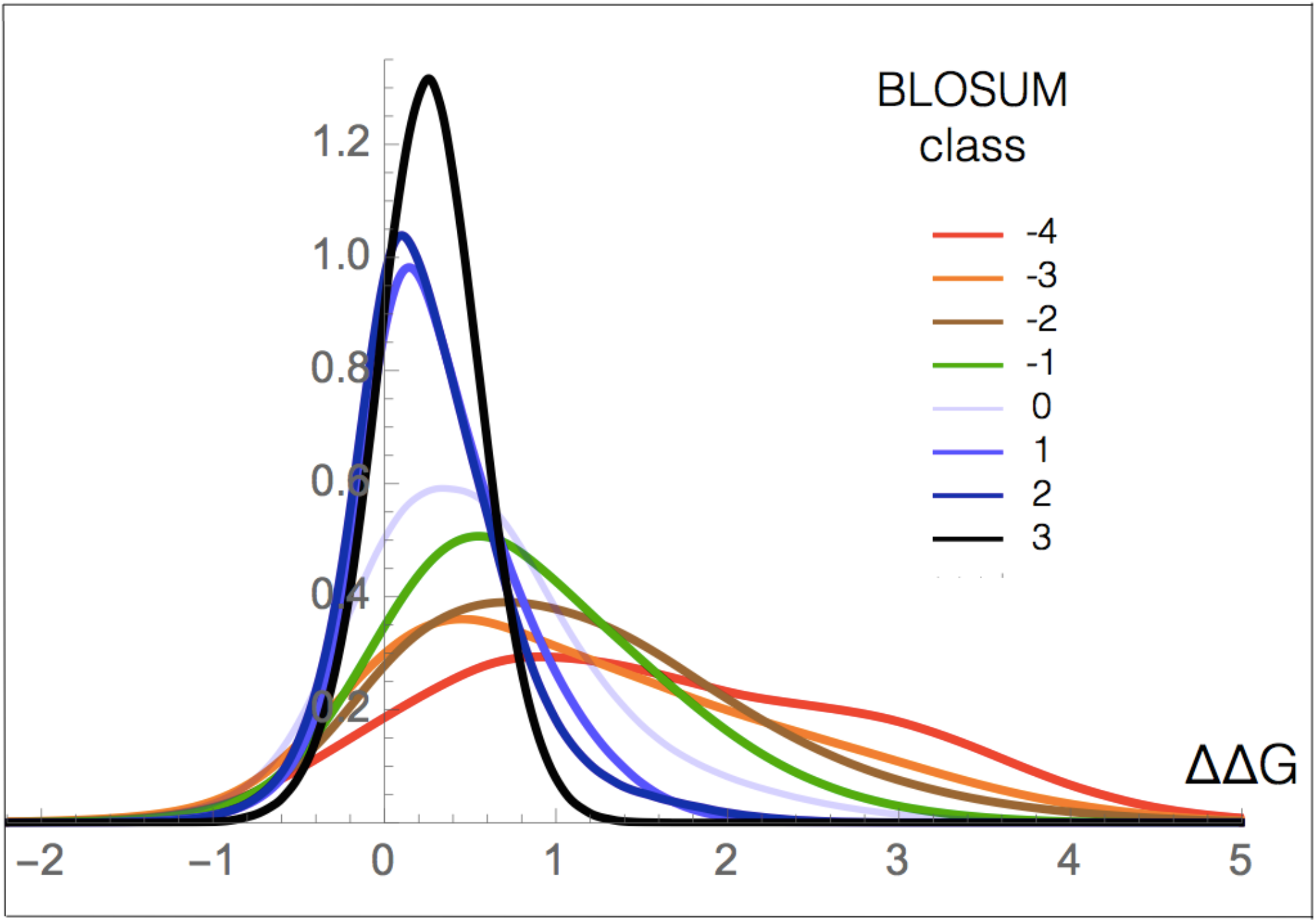
ΔΔ*G* distribution (in kcal/mol) of all *μ*SBSs in 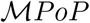 as a function of the BLOSUM62 score. Positive BLOSUM scores indicate more likely amino acid substitutions and negative scores, less likely substitutions.

### 2.6 Structure of the genetic code

We investigated the relation between the mutational robustness and the structure of the standard genetic code. In the codon-to-amino acid mapping, single base substitutions lead to some but not all amino acid mutations. To get them all, the simultaneous substitution of two or three bases has to be considered, which occur at a much lower rate.

We thus compared the mutational ΔΔ*G* profiles of single versus multiple base substitutions (*μ*SBSs versus *μ*MBSs) to better understand the extent to which the standard genetic code is optimized to ensure mutational robustness. Note that we call *μ*MBS, amino acid mutations that cannot be reached by any SBS.

First of all, we found that mutations resulting from single base substitutions are on average less destabilizing than those resulting from multiple base substitutions, for both the core and surface regions (Fig. 4.a-b and Table 3, and Table S1 and Fig. S4 for more detailed RSA dependence). This suggests that the structure of the standard genetic code is optimized, at least partially, for protein mutational robustness through the minimization of the destabilizing impact of random mutations.

**Table 3:** Comparison between the mean ΔΔ*G* values (in kcal/mol) of single and multiple nucleotide substitutions (*μ*SBS and *μ*MBS) in 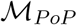 and the fraction of stabilizing, neutral and destabilizing mutations. Core residues have an RSA≤ 20% and surface residues an RSA > 20%.

| Region | $\langle\Delta\Delta G\rangle$ | Stab | Neut | Dest |
| --- | --- | --- | --- | --- |
| μSBS | | | | |
| All | 0.81 | 4% | 41% | 55% |
| Core | 1.09 | 4% | 28% | 68% |
| Surface | 0.49 | 4% | 55% | 41% |
| μMBS | | | | |
| All | 0.97 | 4% | 32% | 64% |
| Core | 1.35 | 4% | 18% | 78% |
| Surface | 0.56 | 5% | 47% | 48% |

**Figure 4:**
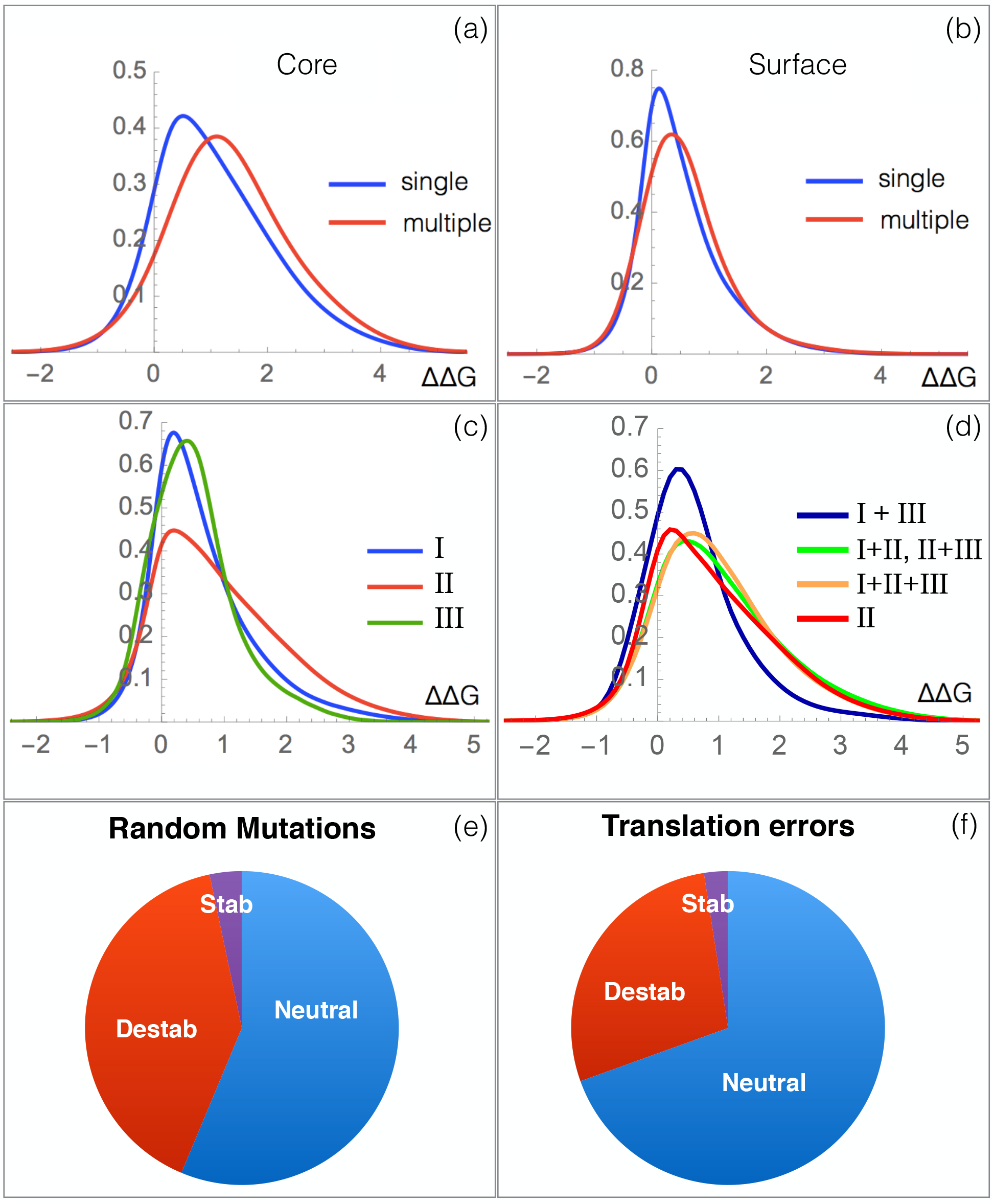
Effects of single and multiple base substitutions and the nucleobase position in the codon. (a)-(d) ΔΔ*G* distribution (in kcal/mol) of amino acid mutations in 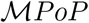. (a) *μ*SBSs and *μ*MBSs in the core (RSA≤ 20%) and (b) on the surface (RSA> 20%); (c) *μ*SBSs resulting from substitutions of bases I, II or III in the codon; (d) *μ*MBSs resulting from simultaneous substitutions of two or three bases in the codon. Ratio of stabilizing, destabilizing and neutral mutations considering random mutations (that occur with equal frequency at each codon position) (e) or translation errors (that occur with different frequency at each codon position) (f). Note that in (e)-(f), the synonymous mutations and mutation degeneracy are included in the computations.

However, a deeper investigation leads to nuance this view. Indeed, there is a large difference according to which bases in the codon are substituted, as seen in Fig. 4.c-d and Table 4. We denote as I, II and III the three bases in the codons.

**Table 4:** Mean ΔΔ*G* (in kcal/mol) for *μ*SBSs from 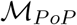 obtained from SBSs at different codon positions (I, II, III). In the lower part of the Table, the mean ΔΔ*G* is computed by considering also the synonymous mutations (with ΔΔ*G* = 0) and the degeneracy (the number of SBSs leading to a *μ*SBS).

| Position | $\langle\Delta\Delta G\rangle$ | Stab | Neut | Dest |
| --- | --- | --- | --- | --- |
| Without synonymous mutations and degeneracy |  |  |  |  |
| I | 0.65 | 3% | 49% | 48% |
| II | 0.91 | 5% | 36% | 59% |
| III | 0.51 | 4% | 50% | 46% |
| With synonymous mutations and degeneracy |  |  |  |  |
| I | 0.64 | 3% | 50% | 47% |
| II | 0.91 | 5% | 36% | 59% |
| III | 0.16 | 2% | 84% | 14% |

Clearly, the substitution of base II in the codon yields the most destabilizing amino acid mutations, on average. At the other extreme, the least destabilizing SBSs involve base III, followed by base I. This is related to the structure of the genetic code and the smallest physicochemical property changes caused by base III substitutions and the largest changes caused by base II substitutions. Again, the trends are more pronounced for core than for surface residues (Table S1 and Fig. S4).

An important result is that we find the same trends with experimental stability values from 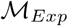 than with computed values from 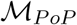, as shown in Table S2.

Moreover, only 14 amino acid mutations are reachable by varying base III, against 64 for base I, and 80 for base II, as can be deduced by looking at the genetic code table (Fig. S1). Thus, not only is base III the most optimized for stability, but it is also the base that leads to the lowest number of non-synonymous mutations. Base II is the least optimized for stability and moreover leads to the highest number of non-synonymous mutations.

As a consequence, the difference between the three base substitutions is even clearer when including the synonymous mutations in the estimation of the mean ΔΔ*G*, which consist of base substitutions that lead to the same amino acid and thus to ΔΔ*G* values equal to zero. We have in that case also to count the degeneracy, that is the number of different base substitutions that yields the same amino acid mutation. The results are shown in Table 4: the mean ΔΔ*G* is lowest for base III (0.16 kcal/mol), medium for base I (0.64 kcal/mol) and highest for base II (0.91 kcal/mol). Analogous differences can be observed at any values of the solvent accessibility and become even more important in the core while decrease at the surface (Table S1).

So there seems to be a stronger positive selection pressure on base I and even more on base III, whereas base II appears much more constrained across evolution. This has sometimes been related to the origin of the genetic code and considered as a by-product of the expansion of the primitive code through the diversification of the amino acids repertoire [18, 47]. However, another interpretation is more straightforward in the present context: our results are related to the codon-anticodon pairing and mispairing in the translation process. Indeed, transfer RNA reads with much higher accuracy base II in the codon than base I and even more, than base III [40, 11].

Our results can thus be taken to mean that natural selection, through targeted adaptation of the standard genetic code, primarily favors an increased translation accuracy, rather than the minimization of the impact of random mutations.

This interpretation is supported by the finding of a high anticorrelation between the mean ΔΔ*G* per position in the codon and the frequency of the translation error at these positions; these frequencies are equal to (31.3%, 6.2%, 62.5%) [36]. Indeed, the Pearson’s linear correlation coefficient is almost perfect: *r* = −0.996 (P-value≃ 0.05).

We also compared the impact of single and multiple nucleotide substitutions (Table S1, Figs 4.c-d and S4). We found that the ΔΔ*G* profile obtained from *μ*MBSs of the two bases I+III are less destabilizing than base II *μ*SBSs, and only slightly more destabilizing than base I or base III *μ*SBSs. Furthermore, the ΔΔ*G* profile of base II *μ*SBSs strongly resembles the profiles of bases I+II and bases II+III *μ*MBSs.

In summary, we have the following increased destabilization ranking: III, I, I+III, II, II+III, I+II, I+II+III. The comparison of these results with the frequency of translation errors yields a very interesting result that further confirms our hypotheses: the anticorrelation between the mean ΔΔ*G* and the frequency of the translation errors for all these different types of substitutions is extremely high *r* = −0.951 (P-value< 0.001).

Finally, we computed the fraction of stabilizing, destabilizing and neutral mutations according to whether they result from random mutations or from errors in translation. In the latter case, the frequencies of the translation errors at the three positions in the codon must be taken into account. As shown in Fig 4.e-f, a much larger number of neutral mutations and a reduced fraction of destabilizing mutations are found if we consider translation errors. This trend is even more pronounced in the core, as seen in Fig. S5.

This result signals a better optimization of the standard genetic code for minimizing the consequences of errors in translation. It is also optimized to minimize the effects of random mutations in the DNA, but to a lesser extent. Indeed, random mutations occur with equal frequency at the three codon positions.

The error rates are known to be of the order of 10^−8^ in genome replication and of 10^−5^ in transcription. Instead, the error rate in protein synthesis is higher with a value of about 10^−4^. This suggests that the mRNA translation process is the real bottleneck in proteome accuracy maintenance [57] and explains our finding that the standard genetic code evolved to primarily favor robustness against mutations caused by defaults in the translation machinery.

### 2.7 Nucleobase composition

Let us now study the mutational robustness as a function of the nucleobase composition of the mRNA sequence, which is often biased and varies from GC- to AT-rich. This composition influences the amino acid content of the encoded protein, as well as protein stability and evolution [56, 24, 25], but the magnitude of this effect is still debated [17].

The mutational robustness was estimated from the mean ΔΔ*G* of *μ*SBSs resulting from the substitution of one of the four nucleobases, independently of their position in the codon (Fig. 5.a and Table S3). We observed that substitutions of A yields the most robust amino acid mutations and substitutions of T the least robust mutations. C and G show similar intermediate behaviors. The same trends are observed for experimental mutations from 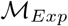 (Table S4).

**Figure 5:**
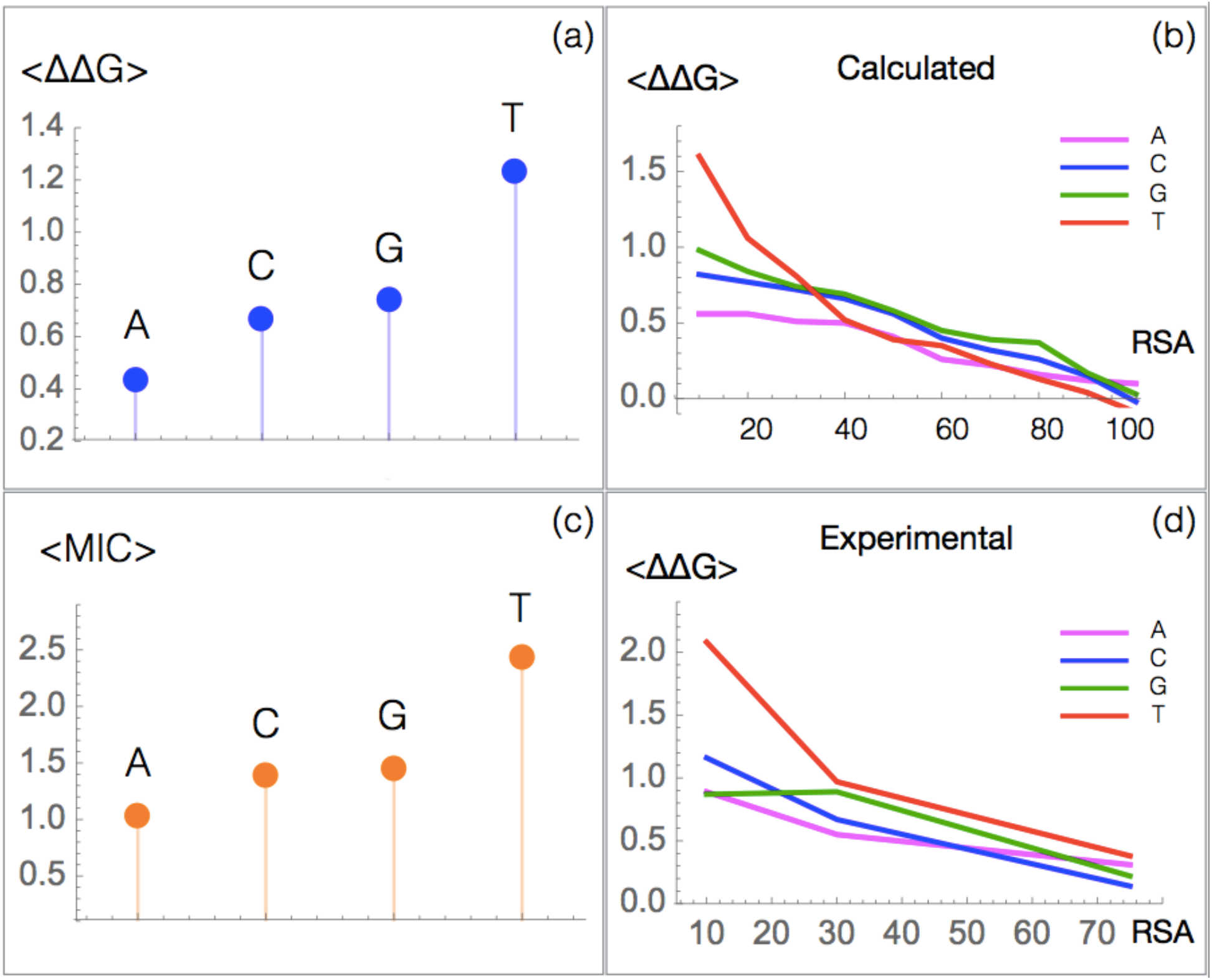
Mean ΔΔ*G* (in kcal/mol) and fitness of amino acid mutations caused by the substitution of each of the four nucleobases. (a) Computed ⟨ΔΔ*G*⟩ of SBSs in the 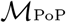 set; (b) Computed ⟨ΔΔ*G*⟩ of SBSs as a function of the residue solvent accessibility (RSA) in the 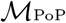 set; (c) Fitness score of mutations measured via the minimum inhibitory concentration (MIC) to *β*-lactam amoxicillin [45]; (d) Computed ⟨ΔΔ*G*⟩ of SBSs as a function of the RSA in the 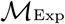 set. Note that here only three RSA bin (0% −20%, 20%-50%, 50%-100%) are considered due to the limited number of entries.

The low robustness of T is not surprising as it is the least frequently mutated base [17, 58] and there is thus a low evolutionary pressure acting on it. In contrast, the high robustness of A is *a priori* surprising since it is usually less frequently mutated than G and C bases.

In fact, the differences observed between the four bases are mainly observed in the protein core. This can be clearly seen from Figs 5.b and d, where the mean ΔΔ*G* as a function of the solvent accessibility is plotted for each kind of wild-type nucleobase, for computed and experimental ΔΔ*G*s.

We can thus hypothesize that these difference are linked to the hydrophobicity of the encoded amino acids. This is indeed the case: the T content of the codons is correlated with the hydrophobicity of the encoded amino acids (*r*=0.55 using the hydrophobicity scale of Kyte-Doolittle, P-value < 10^−7^), the A content is anticorrelated with it (*r*=−0.28, P-value < 0.005). No correlation is observed for C and G.

Thus, T-containing codons code preferentially for hydrophobic amino acids, and their mutations in the core induce on average strong destabilization. In contrast, A-containing codons tend to encode polar amino acids, and their mutations in the core are often neutral or stabilizing. This explains the observed mutational robustness profile upon specific base substitutions and the absence of link with the rate of SBSs.

To better assess our observations, we compared them with the mutagenesis data from 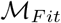. We found a nice agreement between our stability predictions (Fig. 5.a) and the experimental fitness data of TEM-1 *β*-lactamase [45] as measured via the minimum inhibitory concentration (MIC) to *β*-lactam amoxicillin (Fig. 5.c). A similar agreement was found with the other fitness data from 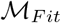 (Fig S6).

### 2.8 Transition:transversion bias

Transitions are substitutions that interchange purines (A ↔ G) or pyrimidines (C ↔ T), whereas transversions interchange purines and pyrimidines (C,T ↔ A,G). Transitions are known to be from 2 to 5 times more frequent than transversions [50, 67], an observation called the transition:transversion bias. However, the origin of this bias is a longstanding problem in molecular evolution.

Recently, the relationship between this bias and the fitness score was analyzed on a set of about 1,200 mutations, in which a probability of 53% was found for the transitions to be fitter than the transversions [67]. However, this tiny difference cannot justify the large bias observed in evolutionary investigations, and thus essentially discard a selection effect as main explanation.

In another recent study [55], transitions were seen to be significantly less detrimental than transversions in deep mutagenesis scanning experiments on the influenza and HIV viruses. This suggests instead that the selective hypothesis cannot be totally ruled out, but that it could contribute, together with other mutational biases, to explain the observed transition:transversion substitution rate.

Our results are basically in agreement with the first aforementioned analysis. Indeed, we found the transitions to be slightly more robust than transversions, with a mean ⟨ΔΔ*G*⟩ of 0.51 and 0.60 kcal/mol, respectively, when considering the mutation degeneracy and the synonymous mutations. However, this free energy difference is too small to explain the large bias observed.

Note that, if only the non-synonymous mutations are included in the ⟨ΔΔ*G*⟩ computation, the opposite trend is observed, both using computed and experimental stability data (Tables 6, S5 and S6). This is due to the fact that transitions are enriched in synonymous mutations.

**Table 5:**
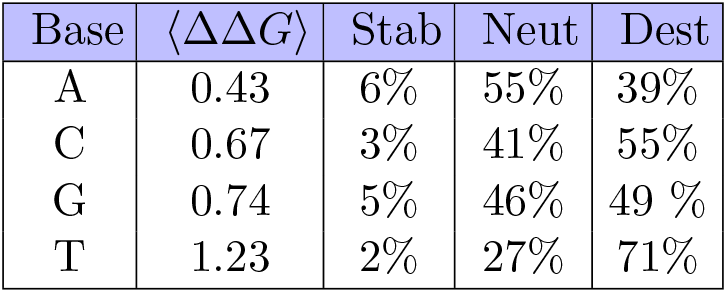
Mean ΔΔ*G* (in kcal/mol) and fraction of stabilizing, neutral and destabilizing *μ*SBSs from 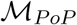 which result from the substitution of one of the four nucleobases.

| Base | $\langle\Delta\Delta G\rangle$ | Stab | Neut | Dest |
| --- | --- | --- | --- | --- |
| A | 0.43 | 6% | 55% | 39% |
| C | 0.67 | 3% | 41% | 55% |
| G | 0.74 | 5% | 46% | 49 % |
| T | 1.23 | 2% | 27% | 71% |

**Table 6:** Mean ΔΔ*G* (in kcal/mol) for *μ*SBSs from 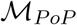 obtained from transitions and transversions. In the upper part of the Table, the mean ΔΔ*G* includes the synonymous mutations (with ΔΔ*G* = 0), while the lower part is without them.

| Mutation | $\langle\Delta\Delta G\rangle$ | Stab | Neut | Dest |
| --- | --- | --- | --- | --- |
| With synonymous mutations |  |  |  |  |
| Transitions | 0.51 | 2% | 63% | 35% |
| Transversions | 0.60 | 4% | 53% | 43% |
| Without synonymous mutations |  |  |  |  |
| Transitions | 0.79 | 3% | 44% | 53% |
| Transversions | 0.73 | 5% | 43% | 52% |

### 2.9 Codon usage

The understanding of the codon usage and its evolution are strongly debated in the molecular evolution field. Indeed, the codon usage is intrinsically connected with a wide range of factors whose contributions are difficult to disentangle [65]. For example, relations of codon usage with tRNA abundance, translation elongation rate, protein expression levels and stability of mRNA secondary structure have been observed, which suggests an explanation in terms of selection for translation efficiency [19, 71, 3].

Another interesting hypothesis is that codon usage is shaped to minimize errors at the protein level. This adaptive hypothesis suggests that a selective pressure for mutational robustness acts on codon usage to reduce the deleterious impacts of genetic variants [5, 4, 6, 2, 28, 79]. In [52], the comparison between wild type and engineered capsid poliovirus, in which synonymous mutations are introduced, suggests that the former has a higher mutational robustness than the latter, and thus that codon choice is directly connected to robustness.

Codon usage could also be related to protein evolvability, since synonymous codons allow the exploration of different evolutionary pathways displaying different sets of proximal amino acid mutations [16].

In order to deepen the hypothesis of the role of the codon usage in minimizing errors at the protein level, we compared the mutational robustness of proteins when using the actual codon or synonymous codons. More specifically, we analyzed how the ⟨ΔΔ*G*⟩ that results from random mutations or translation errors differs according to the codon usage. We also analyzed the ⟨ΔΔ*G*⟩ at each codon position to study a possible position-dependent codon selection.

The difference in ⟨ΔΔ*G*⟩ when using the actual or a synonymous codon is reported in Tables 7 and S7. We observe that the used codons are in general more robust than synonymous ones. The difference can amount to more than 20%. This appears to be true for mutations inserted at each of the three positions in the codon, although to a different extent, and both for random mutations, which do not distinguish between the positions in the codon, and for translation errors, in which the error rate depends on the position.

**Table 7:** Difference between ⟨ΔΔ*G*⟩ for *μ*SBSs in 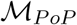 reached from synonymous codons (syn) or from the wild-type codon (used), according to the position of the substituted base in the codon (I, II and III), and according to whether the position-dependent frequency of translation errors is taken into account (translation) or not (random).

| | $\frac{\langle \Delta \Delta G^{syn} \rangle - \langle \Delta \Delta G^{used} \rangle}{\langle \Delta \Delta G^{used} \rangle}$ | | |
| --- | --- | --- | --- |
|  | All | Core | Surface |
| I | 6% | -4% | 24% |
| II | 12% | 7% | 12% |
| III | 21% | 14% | 20% |
| Random | 13% | 6% | 19% |
| Translation | 16% | 8% | 21% |

Interestingly, we found that this effect is smaller in the core and bigger on the surface (Fig. 6). This could be related to the fact that surface residues evolve faster than those in the core. Evolution has thus had the opportunity to better optimize the surface than the core, and this could explain why the chosen codons are more robust in this region. This general trend is independent of the codon position, as seen in Tables 7 and S7, and is also seen for experimentally characterized mutations (Table S8).

**Figure 6:**
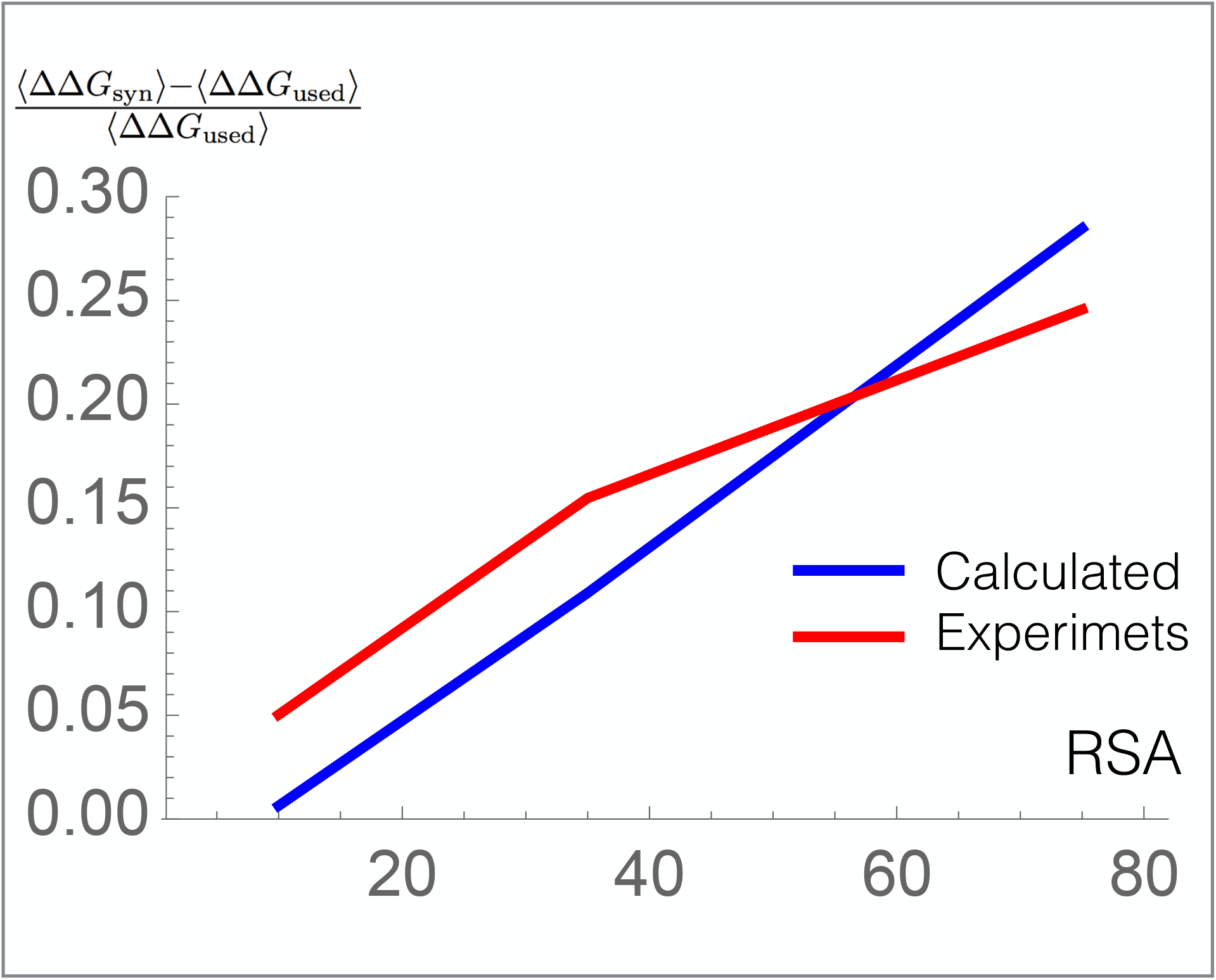
Difference between ⟨ΔΔ*G*⟩ for *μ*SBSs in 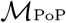 and 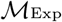 reached from synonymous codons (syn) or from the used codon (used) as a function of the RSA.

Another interesting result is that the codon choice seems to minimize slightly more the effects of translation errors than the impact of random mutations.

We also analyzed the C+G-content in the codons, which is a simple factor related to mRNA stability and translation efficiency [48]. We found that the difference in CG content between used and synonymous codon is equal to be about −4%. This indicates that codons with higher CG content have a slightly higher chance to be used and is probably due to the fact that this choice improves mRNA stability.

### 2.10 Codon usage bias

Some codons occur much more frequent than others, and this effect, known as the codon usage bias, strongly depends on the host organism [8]. This bias has been related to the tRNA pool in the organisms; indeed a correlation between the codon frequency and the concentration of tRNAs with the complementary anticodons has been found in many genomes. This correlation could contribute to the efficiency of the translational process by tuning the elongation rate [61, 41].

We analyzed here whether there is a link between codon choice, codon usage bias and mutational robustness. More precisely, we investigated if the used codon is better optimized for mutational robustness than synonymous codons in the biased or unbiased subsets of codons.

To explore this question, we retrieved the codon usage frequency tables [7] of the host organisms of the proteins from the 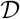 structurome set, and defined codons as biased if their frequency deviates by more than 12.5% from equiprobability [7]. We then compared the ⟨ΔΔ*G*⟩ of *μ*SBSs reached from synonymous and used codons, according to whether these codons are biased or not in the protein’s host organism.

For unbiased codons, the wild-type and synonymous codons appear to have the same mutational robustness (Table 8). In contrast, for biased codons, the used codons appear to be more robust than the synonymous ones.

**Table 8:** Comparison of the mean mutational robustness of used and synonymous codons for *μ*SBSs in 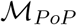 for biased and unbiased codons.

| | $\frac{\langle \Delta \Delta G^{syn} \rangle - \langle \Delta \Delta G^{used} \rangle}{\langle \Delta \Delta G^{used} \rangle}$ | |
| --- | --- | --- |
|  | Biased | Unbiased |
| Random | 16% | 0% |
| Translation | 19% | 1% |

This interesting result suggests that the codon usage bias is not only related to the optimization of the translation efficiency, but also to increase the mutational robustness. It points out the non-trivial role of the selection for error minimization at the protein level in shaping the codon usage, in agreement with an adaptionist hypothesis [5]. Here, for the first time, we quantified these effects that certainly play an important role in the highly complex interdependence with other factors, such as translation elongation speed, initiation efficiency and mRNA stability. These interrelationships need to be further explored.

## 3 Conclusion

The mutational robustness of proteomes and its adaptation across natural evolution are key questions in protein science. Answering these would be proficuous not only for fundamental understanding but also for a wide range of biotechnological and biopharmaceutical applications. To deepen this issue, we investigated here how a series of factors influences protein mutational robustness through large-scale *in-silico* deep-mutagenesis scanning experiments and the analysis of experimental mutagenesis data.

A first point to emphasize is that, whenever the amount of experimental data is sufficient, experimental and computed results largely coincide. This strongly supports the accuracy and unbiased nature of our predictions and the validity of our structurome-scale approach.

Our results can be summarized as follows:

- **Amino acid level.** We quantified the crucial effects of residue RSA and protein length on the folding free energy changes upon mutations, and compared these with previous estimations of evolutionary rates. We found that core residues are much less robust on average and evolve more slowly than mutations on the surface, as they are more constrained. Moreover, short proteins have a less robust core and a more robust surface than longer proteins, as they have larger proportions of buried hydrophobic residues and of exposed functional residues. We also showed that the fraction of stabilizing mutations amount to about 4%, both in the core and on the surface, whereas the fraction of neutral and destabilizing mutations is higher on the surface and in the core, respectively. We found a very nice agreement between these quantities and the fractions of beneficial, neutral and deleterious mutations experimentally estimated in a series of mutagenesis studies. This supports the pivotal role of thermodynamic stability in the fitness cost of mutations [77]. Finally, we found a very high correlation between the mean frequency of mutations across evolution, characterized by the BLOSUM62 matrix, and their mutational robustness: rare mutations are on average more destabilizing that frequent ones.
- **Standard genetic code.** Our analyses at the codon level contributed to get a clearer picture of the impacts of the evolutionary pressure on the standard genetic code. We found that single base substitutions are on average less destabilizing for the protein than multiple base substitutions, which occur at a much lower rate. Our first conclusion is thus that the standard genetic code evolved to minimize the errors of random mutations and to preserve the genome information at all stages, from DNA replication and transcription to mRNA translation and protein synthesis. Interestingly, not all bases in the codon are optimized in the same way. Base II is less robust than base I, which in turn is less robust than base III. This is related to the fact that the translation errors are lowest for base II and highest for base III. Moreover, a better robustness for the joint substitution of bases I and III than for single base II substitutions is found. Strikingly, the linear correlation coefficient between the mean ΔΔ*G* caused by the substitution of specific single and multiple bases and the frequency of translation errors is almost equal to −1. We thus conclude that the genetic code is primarily optimized to limit the mRNA translation errors rather than random mutations. Given that the error rates are higher on translation than on transcription and replication, the minimization of translation errors can be seen as a global optimization of the genetic material encoding.
- **Codon usage and codon usage bias.** We compared the mean change in stability upon mutations when using the wild-type codon or a synonymous codon, and found that the former is on average more robust than the latter, in the sense that SBSs of the wild-type codon yield less destabilizing amino amino acid mutations. The codon is thus selected, at least partly, to minimize the effect of both transcriptional and translational errors. Note that our results show that the codon usage is partially optimized for the precision of translation, but do not exclude that it may also be partially optimized for translation efficiency and for mRNA stability [19, 71, 3, 27]. Interestingly, the codon selection for mutational robustness is stronger at the protein surface, where the mutation rate is higher and thus where natural selection has had more opportunities for optimization. It is moreover stronger for biased than for unbiased codons, suggesting that also the codon usage bias could be partly due to mutational robustness optimization.

We would like to underline that the use of 3D structural information is a fundamental piece in our analyses, which allowed us to gain a deeper understanding of the link between thermodynamic constraints and natural evolution. We believe that this is a general trend, and that the integration of structure and sequence data is needed to further improve our understanding of the evolutionary mechanisms and how the biophysical features shape and are shaped by them.

Different questions still need to be addressed. A first issue is the origin of the mutational robustness and whether it can be considered as an emergent property or a property that depends on an intricate combination of factors, some of which are analyzed in this paper [32]. Other biophysical features such as protein dynamics, conformational disorder or environmental variables such as the organism growth temperature should be explored to better understand the mutational robustness and its highly complex dependencies.

The intricate relation between translation accuracy and efficiency in the codon choice, which involve mutational robustness, protein folding rate, mRNA stability and tRNA abundance, is still an open question. Our study suggests that mutational robustness is an important factor that has to be taken in consideration, but its magnitude with respect to the other contributions need to be clarified.

## 4 Methods

### 4.1 Protein structurome

The non-redundant set 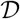 of protein structures analyzed here, which represent the structurome, was obtained by following steps, schematically depicted in Fig. S7:

1. We used the PISCES protein culling server [74] to get the subset of proteins from the Protein Data Bank (PDB) [9] which have an experimental X-ray structure of at most 2.5Å resolution, and share less than 95% pairwise sequence identity.
2. We filtered out the membrane proteins, viral capsid proteins and antibodies on the basis of PDB annotations. The first series of proteins is overlooked because the PoPMuSiC^*sym*^ predictor is applicable to globular proteins only, the second series because they form very large oligomeric assemblies, and the last because antibodies have variable sequences and the mutations in their complementarity determining regions have a clear functional role. We obtained in this way a uniform set of globular proteins.
3. For each protein entry, we retrieved the DNA sequence from the EMBL webserver [49].
4. To check the protein-DNA mapping, we aligned the translated DNA sequences with the protein sequences from the PDB using CLUSTALW [68]. We kept only the DNA sequences which yield at least 95% sequence identity with the PDB sequences.

With this procedure, we obtained 21,540 X-ray structures amounting to 5,368,279 residues in total.

### 4.2 Large-scale *in silico* mutagenesis experiments

We estimated the folding free energy changes (ΔΔ*G*) caused by all possible single-site mutations introduced in all collected structures, using the unbiased version of our in-house predictor PoPMuSiC, called PoPMuSiC^*sym*^ [60, 59]. The set of mutations so obtained is called 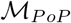. It contains 101,997,301 mutations among which 100,149,646 have a known wild-type codon.

The model structure of PoPMuSiC^*sym*^ consists of a linear combination of energy terms estimated using different types of statistical potentials. These have been derived from frequencies of sequence-structure associations in sets of protein X-ray structures, transformed into folding free energies using the Boltzmann law. The coefficients of the linear combinations are sigmoid functions of the RSA of the mutated residues.

The model structure of PoPMuSiC^*sym*^ has been designed to avoid the biases toward destabilizing mutations, which we have shown to be present in the original PoPMuSiC version [59]. The results are thus expected to be more reliable when used in large-scale *in silico* mutagenesis experiments. Indeed, a similar analysis using the biased PoPMuSiC version confirms that PoPMuSiC^*sym*^ yields results that are much closer to the experimental data.

We refer to [60] for further details about the PoPMuSiC^*sym*^ predictor.

### 4.3 Experimentally characterized stability changes

We also considered the set of 2,648 mutations of which the ΔΔ*G* folding free energy change upon single-site mutations has been experimentally measured. This set, that we call 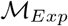, was manually curated as described in [21] and was further annotated according to the previously described pipeline. It has been used to train the PoPMuSiC predictors. The list of mutations of 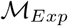 can be found in the supplementary material of [21].

## Acknowledgments

We thank the FNRS - Fund for Scientific Research for its financial support through a PDR research project. FP and MR are FNRS postdoctoral researcher and research director, respectively, and MS benefits from a FRIA PhD grant.

## References

[1] Achoch, M., Dorantes-Gilardi, R., Wymant, C., Feverati, G., Salamatian, K., Vuillon, L., and Lesieur, C. 2016. Protein structural robustness to mutations: an in silico investigation. Phys Chem Chem Phys, 18(20): 13770–13780.

[2] Akashi, H. 1994. Synonymous codon usage in Drosophila melanogaster: natural selection and translational accuracy. Genetics, 136(3): 927–935.

[3] Akashi, H. and Eyre-Walker, A. 1998. Translational selection and molecular evolution. Curr. Opin. Genet. Dev., 8(6): 688–693.

[4] Archetti, M. 2004. Selection on codon usage for error minimization at the protein level. J. Mol. Evol., 59(3): 400–415.

[5] Archetti, M. 2006. Genetic robustness and selection at the protein level for synonymous codons. J. Evol. Biol., 19(2): 353–365.

[6] Archetti, M. 2009. Genetic robustness at the codon level as a measure of selection. Gene, 443(1-2): 64–69.

[7] Athey, J., Alexaki, A., Osipova, E., Rostovtsev, A., Santana-Quintero, L. V., Katneni, U., Simonyan, V., and Kimchi-Sarfaty, C. 2017. A new and updated resource for codon usage tables. BMC Bioinformatics, 18(1): 391.

[8] Behura, S. K. and Severson, D. W. 2013. Codon usage bias: causative factors, quantification methods and genome-wide patterns: with emphasis on insect genomes. Biological Reviews, 88(1): 49–61.

[9] Berman, H. M., Westbrook, J., Feng, Z., Gilliland, G., Bhat, T. N., Weissig, H., Shindyalov, I. N., and Bourne, P. E. 2000. The Protein Data Bank. Nucleic Acids Res., 28(1): 235–242.

[10] Besenmatter, W., Kast, P., and Hilvert, D. 2007. Relative tolerance of mesostable and thermostable protein homologs to extensive mutation. Proteins, 66(2): 500–506.

[11] Blazej, P., Wnetrzak, M., Mackiewicz, D., and Mackiewicz, P. 2018. Correction: Optimization of the standard genetic code according to three codon positions using an evolutionary algorithm. PLoS ONE, 13(10): e0205450.

[12] Bloom, J. D., Silberg, J. J., Wilke, C. O., Drummond, D. A., Adami, C., and Arnold, F. H. 2005. Thermodynamic prediction of protein neutrality. Proc. Natl. Acad. Sci. U.S.A., 102(3): 606–611.

[13] Bloom, J. D., Labthavikul, S. T., Otey, C. R., and Arnold, F. H. 2006. Protein stability promotes evolvability. Proc. Natl. Acad. Sci. U.S.A., 103(15): 5869–5874.

[14] Bloom, J. D., Lu, Z., Chen, D., Raval, A., Venturelli, O. S., and Arnold, F. H. 2007a. Evolution favors protein mutational robustness in sufficiently large populations. BMC Biol., 5: 29.

[15] Bloom, J. D., Raval, A., and Wilke, C. O. 2007b. Thermodynamics of neutral protein evolution. Genetics, 175(1): 255–266.

[16] Cambray, G. and Mazel, D. 2008. Synonymous genes explore different evolutionary landscapes. PLoS Genet., 4(11): e1000256.

[17] Chen, S. L., Lee, W., Hottes, A. K., Shapiro, L., and McAdams, H. H. 2004. Codon usage between genomes is constrained by genome-wide mutational processes. Proc. Natl. Acad. Sci. U.S.A., 101(10): 3480–3485.

[18] Chiusano, M. L., Alvarez-Valin, F., Di Giulio, M., D’Onofrio, G., Ammirato, G., Colonna, G., and Bernardi, G. 2000. Second codon positions of genes and the secondary structures of proteins. Relationships and implications for the origin of the genetic code. Gene, 261(1): 63–69.

[19] Coleman, J. R., Papamichail, D., Skiena, S., Futcher, B., Wimmer, E., and Mueller, S. 2008. Virus attenuation by genome-scale changes in codon pair bias. Science, 320(5884): 1784–1787.

[20] De Laet, M., Gilis, D., and Rooman, M. 2016. Stability strengths and weaknesses in protein structures detected by statistical potentials: Application to bovine seminal ribonuclease. Biophysical Journal, 84: 143–158.

[21] Dehouck, Y., Grosfils, A., Folch, B., Gilis, D., Bogaerts, P., and Rooman, M. 2009. Fast and accurate predictions of protein stability changes upon mutations using statistical potentials and neural networks: PoPMuSiC-2.0. Bioinformatics, 25(19): 2537–2543.

[22] Dehouck, Y., Gilis, D., and Rooman, M. 2014. Database-derived potentials dependent on protein size for in silico folding and design. Biophysical Journal, 87: 171–181.

[23] Di Giulio, M. and Medugno, M. 1999. Physicochemical optimization in the genetic code origin as the number of codified amino acids increases. J. Mol. Evol., 49(1): 1–10.

[24] D’Onofrio, G., Mouchiroud, D., Aissani, B., Gautier, C., and Bernardi, G. 1991. Correlations between the compositional properties of human genes, codon usage, and amino acid composition of proteins. J. Mol. Evol., 32(6): 504–510.

[25] D’Onofrio, G., Jabbari, K., Musto, H., and Bernardi, G. 1999. The correlation of protein hydropathy with the base composition of coding sequences. Gene, 238(1): 3–14.

[26] Draghi, J., Parsons, T., Wagner, G., and Plotkin, J. 2010. Mutational robustness can facilitate adaptation. Nature, 463: 353–355.

[27] Drummond, D. A. and Wilke, C. O. 2008. Mistranslation-induced protein misfolding as a dominant constraint on coding-sequence evolution. Cell, 134(2): 341 – 352.

[28] Drummond, D. A. and Wilke, C. O. 2009. The evolutionary consequences of erroneous protein synthesis. Nat. Rev. Genet., 10(10): 715–724.

[29] Echave, J., Jackson, E. L., and Wilke, C. O. 2015. Relationship between protein thermodynamic constraints and variation of evolutionary rates among sites. Physical Biology, 12(2): 025002.

[30] Echave, J., Spielman, S. J., and Wilke, C. O. 2016. Causes of evolutionary rate variation among protein sites. Nature Reviews Genetics, 17(2): 109–121.

[31] Epstein, C. J. 1966. Role of the amino-acid “code” and of selection for conformation in the evolution of proteins. Nature, 210(5031): 25–28.

[32] Fares, M. A. 2015. The origins of mutational robustness. Trends Genet., 31(7): 373–381.

[33] Faure, G. and Koonin, E. V. 2015. Universal distribution of mutational effects on protein stability, uncoupling of protein robustness from sequence evolution and distinct evolutionary modes of prokaryotic and eukaryotic proteins. Phys Biol, 12(3): 035001.

[34] Franzosa, E. A. and Xia, Y. 2009. Structural determinants of protein evolution are context-sensitive at the residue level. Mol. Biol. Evol., 26(10): 2387–2395.

[35] Franzosa, E. A. and Xia, Y. 2012. Independent effects of protein core size and expression on residue-level structure-evolution relationships. PLoS ONE, 7(10): e46602.

[36] Freeland, S. J. and Hurst, L. D. 1998. The genetic code is one in a million. J. Mol. Evol., 47(3): 238–248.

[37] Freiberger, M., Guzovsky, A., Wolynes, P., Parra, R., and D.U., F. 2019. Local frustration around enzyme active sites. Proc Natl Acad Sci U S A, 116: 4037–4043.

[38] Gilis, D., Massar, S., Cerf, N. J., and Rooman, M. 2001. Optimality of the genetic code with respect to protein stability and amino-acid frequencies. Genome Biol., 2(11): RE-SEARCH0049.

[39] Goldberg, A. L. and Wittes, R. E. 1966. Genetic code: aspects of organization. Science, 153(3734): 420–424.

[40] Haig, D. and Hurst, L. D. 1991. A quantitative measure of error minimization in the genetic code. J. Mol. Evol., 33(5): 412–417.

[41] Hanson, G. and Coller, J. 2018. Codon optimality, bias and usage in translation and mrna decay. Nature Reviews Molecular Cell Biology, 19(1): 20–30.

[42] Henikoff, S. and Henikoff, J. G. 1992. Amino acid substitution matrices from protein blocks. Proc. Natl. Acad. Sci. U.S.A., 89(22): 10915–10919.

[43] Ikemura, T. 1981. Correlation between the abundance of Escherichia coli transfer RNAs and the occurrence of the respective codons in its protein genes: a proposal for asynonymous codon choice that is optimal for the E. coli translational system. J. Mol. Biol., 151: 389–-409.

[44] Ikemura, T. 1985. Codon usage and tRNA content in unicellularand multicellular organisms. Mol. Biol. Evol., 2: 13–34.

[45] Jacquier, H., Birgy, A., Le Nagard, H., Mechulam, Y., Schmitt, E., Glodt, J., Bercot, B., Petit, E., Poulain, J., Barnaud, G., Gros, P. A., and Tenaillon, O. 2013. Capturing the mutational landscape of the beta-lactamase TEM-1. Proc. Natl. Acad. Sci. U.S.A., 110(32): 13067–13072.

[46] Kimura, M. 1968. Evolutionary rate at the molecular level. Nature, 217(5129): 624–626.

[47] Koonin, E. V. and Novozhilov, A. S. 2009. Origin and evolution of the genetic code: the universal enigma. IUBMB Life, 61(2): 99–111.

[48] Kudla, G., Lipinski, L., Caffin, F., Helwak, A., and Zylicz, M. 2006. High guanine and cytosine content increases mrna levels in mammalian cells. PLOS Biology, 4(6).

[49] Kulikova, T., Akhtar, R., Aldebert, P., Althorpe, N., Andersson, M., Baldwin, A., Bates, K., Bhattacharyya, S., Bower, L., Browne, P., Castro, M., Cochrane, G., Duggan, K., Eberhardt, R., Faruque, N., Hoad, G., Kanz, C., Lee, C., Leinonen, R., Lin, Q., Lombard, V., Lopez, R., Lorenc, D., McWilliam, H., Mukherjee, G., Nardone, F., Pastor, M. P., Plaister, S., Sobhany, S., Stoehr, P., Vaughan, R., Wu, D., Zhu, W., and Apweiler, R. 2007. EMBL Nucleotide Sequence Database in 2006. Nucleic Acids Res., 35(Database issue): 16–20.

[50] Kumar, S. 1996. Patterns of nucleotide substitution in mitochondrial protein coding genes of vertebrates. Genetics, 143(1): 537–548.

[51] Lassig, M., Mustonen, V., and Walczak, A. M. 2017. Predicting evolution. Nat Ecol Evol, 1(3): 77.

[52] Lauring, A., Acevedo, A., Cooper, S., and Andino, R. 2012. Codon usage determines the mutational robustness, evolutionary capacity, and virulence of an rna virus. Cell Host & Microbe, 12(5): 623–632.

[53] Lenski, R. E., Barrick, J. E., and Ofria, C. 2006. Balancing robustness and evolvability. PLoS Biol., 4(12): e428.

[54] Lind, P. A., Arvidsson, L., Berg, O. G., and Andersson, D. I. 2017. Variation in Mutational Robustness between Different Proteins and the Predictability of Fitness Effects. Mol. Biol. Evol., 34(2): 408–418.

[55] Lyons, D. M. and Lauring, A. S. 2017. Evidence for the Selective Basis of Transition-to-Transversion Substitution Bias in Two RNA Viruses. Mol. Biol. Evol., 34(12): 3205–3215.

[56] Mendez, R., Fritsche, M., Porto, M., and Bastolla, U. 2010. Mutation bias favors protein folding stability in the evolution of small populations. PLoS Comput. Biol., 6(5): e1000767.

[57] Mohler, K. and Ibba, M. 2017. Translational fidelity and mistranslation in the cellular response to stress. Nature Microbiology, 2(9): 17117.

[58] Pucci, F. and Rooman, M. 2019. Relation between DNA ionization potentials, single base substitutions and pathogenic variants. BMC Genomics, 20(Suppl 8): 551.

[59] Pucci, F., Bernaerts, K., Teheux, F., Gilis, D., and Rooman, M. 2015. Symmetry principles in optimization problems: an application to protein stability prediction. IFAC-PapersOnLine, 48(1): 458–463.

[60] Pucci, F., Bernaerts, K. V., Kwasigroch, J. M., and Rooman, M. 2018. Quantification of biases in predictions of protein stability changes upon mutations. Bioinformatics, 34(21): 3659–3665.

[61] Quax, T. F., Claassens, N., Söll, D., and van?der?Oost, J. 2015. Codon bias as a means to fine-tune gene expression. Molecular Cell, 59(2): 149–161.

[62] Ramsey, D. C., Scherrer, M. P., Zhou, T., and Wilke, C. O. 2011. The relationship between relative solvent accessibility and evolutionary rate in protein evolution. Genetics, 188(2): 479–488.

[63] Scherrer, M. P., Meyer, A. G., and Wilke, C. O. 2012. Modeling coding-sequence evolution within the context of residue solvent accessibility. BMC Evol. Biol., 12: 179.

[64] Serohijos, A. W., Rimas, Z., and Shakhnovich, E. I. 2012. Protein biophysics explains why highly abundant proteins evolve slowly. Cell Rep, 2(2): 249–256.

[65] Shah, P. and Gilchrist, M. A. 2011. Explaining complex codon usage patterns with selection for translational efficiency, mutation bias, and genetic drift. Proc. Natl. Acad. Sci. U.S.A., 108(25): 10231–10236.

[66] Sikosek, T. and Chan, H. S. 2014. Biophysics of protein evolution and evolutionary protein biophysics. J R Soc Interface, 11(100): 20140419.

[67] Stoltzfus, A. and Norris, R. W. 2016. On the Causes of Evolutionary Transition:Transversion Bias. Mol. Biol. Evol., 33(3): 595–602.

[68] Thompson, J. D., Higgins, D. G., and Gibson, T. J. 1994. CLUSTAL W: improving the sensitivity of progressive multiple sequence alignment through sequence weighting, position-specific gap penalties and weight matrix choice. Nucleic Acids Res., 22(22): 4673–4680.

[69] Tokuriki, N. and Tawfik, D. S. 2009a. Protein dynamism and evolvability. Science, 324(5924): 203–207.

[70] Tokuriki, N. and Tawfik, D. S. 2009b. Stability effects of mutations and protein evolvability. Curr. Opin. Struct. Biol., 19(5): 596–604.

[71] Tuller, T., Waldman, Y. Y., Kupiec, M., and Ruppin, E. 2010. Translation efficiency is determined by both codon bias and folding energy. Proc. Natl. Acad. Sci. U.S.A., 107(8): 3645–3650.

[72] van Nimwegen, E., Crutchfield, J. P., and Huynen, M. 1999. Neutral evolution of mutational robustness. Proc. Natl. Acad. Sci. U.S.A., 96(17): 9716–9720.

[73] Wagner, A. 2008. Robustness and evolvability: a paradox resolved. Proceedings of The Royal Society B, 275: 91–100.

[74] Wang, G. and Dunbrack, R. L. 2003. PISCES: a protein sequence culling server. Bioinformatics, 19(12): 1589–1591.

[75] Weile, J., Sun, S., Cote, A. G., Knapp, J., Verby, M., Mellor, J. C., Wu, Y., Pons, C., Wong, C., van Lieshout, N., Yang, F., Tasan, M., Tan, G., Yang, S., Fowler, D. M., Nussbaum, R., Bloom, J. D., Vidal, M., Hill, D. E., Aloy, P., and Roth, F. P. 2017. A framework for exhaustively mapping functional missense variants. Mol. Syst. Biol., 13(12): 957.

[76] Wnętrzak, M., Błażej, P., Mackiewicz, D., and Mackiewicz, P. 2018. The optimality of the standard genetic code assessed by an eight-objective evolutionary algorithm. BMC Evol. Biol., 18(1): 192.

[77] Wylie, C. S. and Shakhnovich, E. I. 2011. A biophysical protein folding model accounts for most mutational fitness effects in viruses. Proc. Natl. Acad. Sci. U.S.A., 108(24): 9916–9921.

[78] Yeh, S. W., Liu, J. W., Yu, S. H., Shih, C. H., Hwang, J. K., and Echave, J. 2014. Site-specific structural constraints on protein sequence evolutionary divergence: local packing density versus solvent exposure. Mol. Biol. Evol., 31(1): 135–139.

[79] Zhou, T., Weems, M., and Wilke, C. O. 2009. Translationally Optimal Codons Associate with Structurally Sensitive Sites in Proteins. Molecular Biology and Evolution, 26(7): 1571–1580.

